# Unraveling the Global Proteome and Phosphoproteome of Prostate Cancer Patient-Derived Xenografts

**DOI:** 10.1101/2023.08.02.551697

**Authors:** Zoi E. Sychev, Abderrahman Day, Hannah E. Bergom, Gabrianne Larson, Atef Ali, Megan Ludwig, Ella Boytim, Ilsa Coleman, Eva Corey, Stephen R. Plymate, Peter S. Nelson, Justin H. Hwang, Justin M. Drake

## Abstract

Resistance to androgen deprivation therapies leads to metastatic castration-resistant prostate cancer (mCRPC) of adenocarcinoma (AdCa) origin that can transform to emergent aggressive variant prostate cancer (AVPC) which has neuroendocrine (NE)-like features. To this end, we used LuCaP patient-derived xenograft (PDX) tumors, clinically relevant models that reflects and retains key features of the tumor from advanced prostate cancer patients. Here we performed proteome and phosphoproteome characterization of 48 LuCaP PDX tumors and identified over 94,000 peptides and 9,700 phosphopeptides corresponding to 7,738 proteins. When we compared 15 NE versus 33 AdCa PDX samples, we identified 309 unique proteins and 476 unique phosphopeptides that were significantly altered and corresponded to proteins that are known to distinguish these two phenotypes. Assessment of protein and RNA concordance from these tumors revealed increased dissonance in transcriptionally regulated proteins in NE and metabolite interconversion enzymes in AdCa.

## Introduction

Prostate cancer (PCa) is the most diagnosed cancer in men in the United States. Early detection via regular screening of serum prostate-specific antigen (PSA) levels have facilitated PCa diagnosis in organ confined tumors before cancer spread ^1^. If a patient presents with aggressive PCa, a classic upfront therapy involves radiation or surgery with androgen-deprivation therapy (ADT) ^2, 3^. While ADT response is effective initially, tumors progress to a more aggressive disease known as metastatic castration-resistance prostate cancer (mCRPC) ^3, 4^. Treatment of mCRPC with adenocarcinoma (AdCa) features consists of hormonal therapies such as enzalutamide, abiraterone acetate, darolutamide, and apalutamide; however, these therapies can often induce novel phenotypes such as aggressive variant prostate cancer (AVPC). AVPC has genetic aberrations, including *PTEN* and *RB1* loss, *TP53* mutations, and diminished AR signaling activity. Several definitions of AVPC have been described including treatment-emergent small cell carcinoma, double-negative prostate cancer, amphicrine, or neuroendocrine prostate cancer (NEPC) ^5, 6^. Emergence of these drug-resistant phenotypes creates a large unmet medical need to identify new protein or phosphoprotein drug targets for potential biomarker and therapeutic development for this subset of CRPC patients.

Analysis of the genomic aberrations has contributed to the understanding of drug resistance mechanisms in PCa. These include mutations and focal amplifications in the androgen receptor (AR), *PIK3CA/B,* fusions in *BRAF/RAF1,* mutations in *APC,* mutations and amplifications in *CTNNB1,* focal homozygous deletions in *ZBTB16/PLZF,* biallelic loss, inactivation and somatic point mutations in *BRCA1* and *BRCA2* and biallelic loss and point mutations in *ATM* ^7^. Additional alteration among other biologically relevant genes includes point mutations of *SPOP, FOXA1*, and *TP53*; *MYC, RB1, PTEN*, and *CHD1* copy number alterations; and *E26* transformation-specific (ETS) fusions, ^8–15^. The identification of such alterations has paved the way to define pathological categorizations of mCRPC between AR+ (AdCa) and AR-(NE) disease states. These two types of metastatic phenotypes show distinct pathological features, which are in most cases consequences of implementation of different treatment modalities with no curative therapies available. In solid tumors, prostate cancer nonsynchronous mutational rate is in the lower 25% quartile compared to esophageal and colorectal tumors which are in the higher 75% quartile ^7, 16–22^. Furthermore, somatic alterations, such as in the PI3K pathway, is present in only 49% (73/150) of the mCRPC afflicted patients, and still nearly half of this population will fail to respond if treated with PI3K inhibitors, ^7, 9, 23^. Overall, this information indicates that the genomic feature of prostate cancer explains some of the tumor progression and therapy responses but dismiss key phenotypic expression from proteins that may be driving the biology and drug resistance.

Several patient-derived xenograft tumor (PDX) models have been developed in prostate cancer reflecting different clinical subtypes including the typical prostate AdCa and the atypical patterns of progression known as AVPC that includes NE tumors. These models have shown to closely reflect the characteristics of the heterogeneity of the patient tumor population, maintaining histopathologic architecture and the genomic footprint of the tumors from which they were derived ^24–28^. The LuCaP series have been extensively characterized, including analyses of genomic alterations, transcriptomic profiles, and single tandem repeats ^29^. However, little is known about the proteomic profiles of these tumors and specifically post-translational modification of these proteins such as phosphorylation.

Recent technological advancements in mass spectrometry (MS) based proteomics have allowed the increase in protein detection, coverage, and quantification ^30, 31^. Here, we used Field Asymmetric Ion Mobility Spectrometry (FAIMS) [31] technology and performed a global proteome and phosphoproteome analysis of the LuCaP PDX models to elucidate proteome-wide signatures and unique activated pathways between AdCa and AVPC (with an emphasis on NEPC). We have developed the largest proteome and phosphoproteome database of prostate cancer PDX models which provides an extensive list of protein targets for drug development, predicted kinase enriched domains, found in blood (plasma), secreted, surface proteins, and biological pathways. Most importantly, we integrated RNA sequencing data from the same PDX models with our proteomic data and evaluated the concordance between protein and mRNA. The results revealed dissonance between protein and mRNA expression of some important biological targets that would have been dismissed if only RNA was analyzed from these tumors. Overall, this large proteomic and phosphoproteomics resource will aid to a further understanding of the underlying cell signaling mechanisms, identification and functional validation of novel drug targets, and future biomarker development in prostate cancer.

## Results

### Proteomics and phosphoproteomics platform analysis

We developed a global systematic approach to evaluate the proteome and phosphoproteome of 48 PDX tumor samples, which includes 6 different AdCa tumors grown in intact mice (AdCa NCR, n=15), 6 AdCa grown in castrated mice (AdCa CR, n=18) and 6 neuroendocrine (NE, n=15) tumors (**Fig 1A-sample collection**). These samples were all processed in parallel on the same day and bottom-up proteomics was performed **(Fig 1A-sample processing).** To evaluate the phosphoproteome, we performed a tyrosine (Y), Serine (S) and Threonine (T) enrichment analysis using a sequential metal oxide affinity chromatography (SMOAC) assay **(Fig 1A-sample processing-II).** In parallel, from the pool of peptides prior to phospho-STY enrichment, we used this fraction and evaluated the overall peptide mix, which we defined as the proteome **(Fig 1A sample processing-I).** The enriched phosphopeptides and total peptides were analyzed using a state-of-art instruments to date, containing a high-field asymmetric waveform ion mobility spectrometry (FAIMS), which is an atmospheric pressure ion mobility technique that separates gas-phase ions by their behavior in strong and weak electric fields. This approach allows better separation and detection of stable peptides (> +2 charge state ions) for confident quantification ^31^ compared to no FAIMS application where the instrument collects +1 charge state ions, which are very unstable and not quantifiable among other advantageous features **(Fig 1A).** We used an in-house proteome and phosphoproteome analysis pipeline ^32^ that includes Maxquant ^33^ for peptide/phosphopeptides searches and data processing **(Supplemental Fig 1.A).** Using a 1% FDR, we identified a total of 94,517 peptides that mapped to 7,738 master proteins at the proteome level and a total of 9,722 phosphopeptides that mapped to 3,759 phosphoproteins **(Fig 1B and 1C)** with a phosphosite probability > 0.75. In combination, we identified 8,612 unique master proteins from these samples, where 32.6% of these proteins overlapped between the proteome and phosphoproteome datasets. After identifying the overall number of master proteins measured between the phosphoproteome and proteome, 9.9 % of the phosphorylated peptides were unique to the phosphoproteome and 57.5% were unique to proteome **(Fig 1D).** This indicates that not all proteins have a phosphorylated peptides and not all the phosphopeptides had a non-phosphorylated protein measured in the proteome.

**Figure 1.**
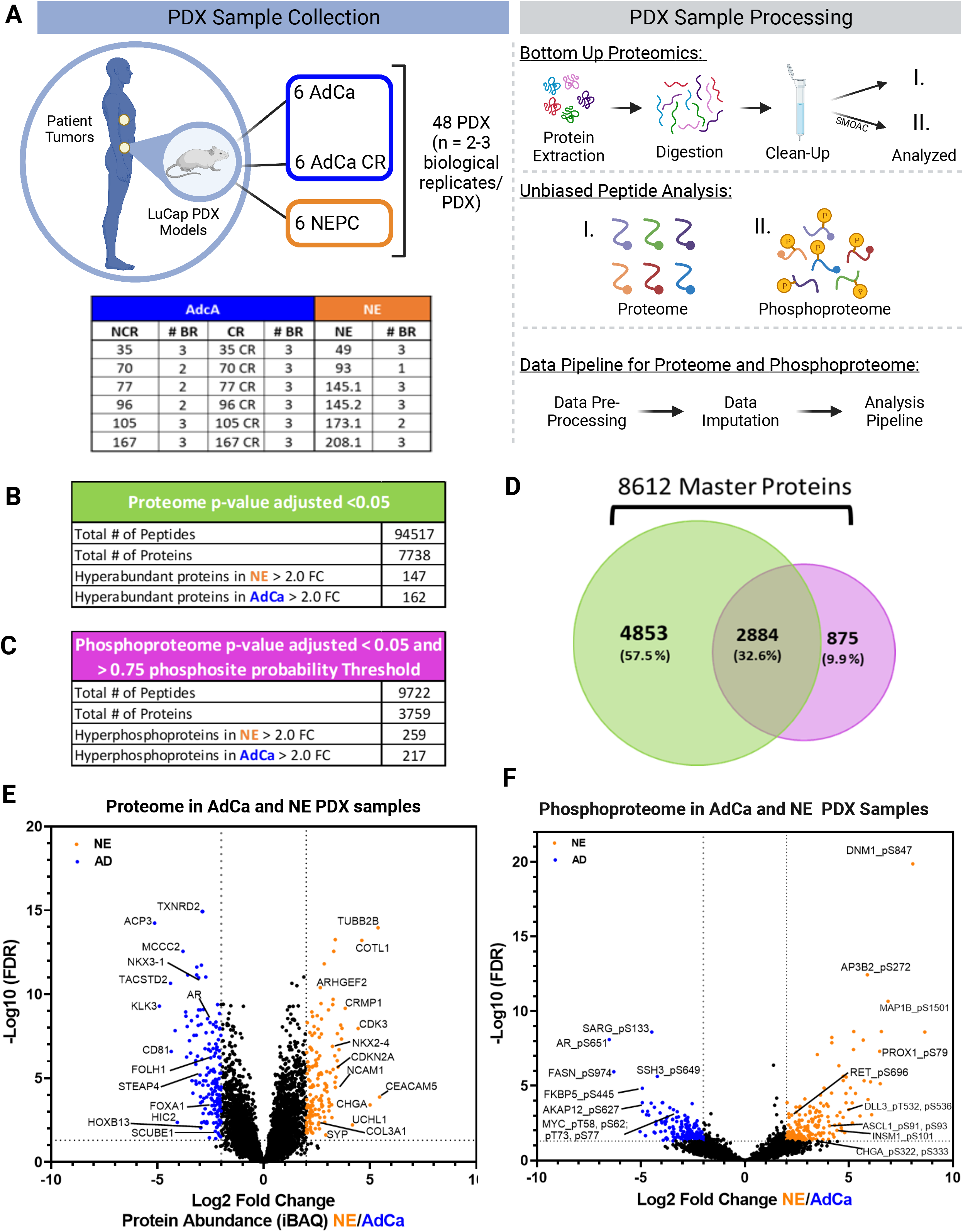
Proteomics and Phosphoproteomics platform analysis. **A.** The LuCaP series of 48 patient derived xenografts (PDX) tumors depicted in the table, where 33 Adenocarcinoma (AdCa) either castrated and non-castrated (all) tumors are shown in dark blue and 15 neuroendocrine prostate cancer (NEPC) tumors are shown in orange, n=2-3 biological replicates (BR). The PDXs were processed by extracting proteins and an enzymatic digestion was performed using Trypsin and LysC. Peptides were purified by reversed-phase chromatography. The final peptide pool was ran as the proteome (I.) and in parallel from this peptide pool a sequential metal oxide affinity chromatography was performed to enrich for phosphorylated Serine, Threonine and Tyrosine (II.). Finally, raw data was searched, processed and analyzed. **B.** Overall proteome results using 1%FDR for protein identification and p-value adjusted < 0.05 log2 fold change significance. **C.** Overall proteome results using 1%FDR for phosphoprotein identification and p-value adjusted < 0.05 log2 fold change significance and >.75 phosphosite probability threshold. **D.** Venn diagram shows the total number of 8612 master proteins identified when both data sets are overlaid. **E.** Volcano plot depicting the intensity based average quantification (iBAQ) enriched in NE and AdCa. **F.** Volcano plot of the phosphoprotein enriched in NE vs AdCa. Grayed line in the x-axis and y-axis are the cut off threshold for NE 2-fold change and for AdCa -2-fold change and p value adjusted to (-log10 FDR), respectively in E-F.

To infer protein abundances, we used intensity-based absolute quantification (iBAQ) ^34^. To identify the significantly altered proteins, we performed a variance stabilization normalization (VSN) ^35^ making the sample variances nondependent from their mean intensities using p-value adjusted > 0.05 and a log 2-fold change. Using this approach, we identified 147 proteins that were hyper-abundant in the NE group and 162 proteins in the AdCa group **(Fig 1E).** Using similar statistical analyses, we identified 259 unique hyperphosphorylated peptides in the NE samples and 217 unique hyperphosphorylated peptides in the AdCa group **(Fig 1F).** We performed several comparisons between AdCa grown in castrated mice (CR) versus AdCa grown in intact mice (NCR), AdCa-CR versus NE, and then AdCa-NCR and AdCa-CR versus NE **(supplemental Table 1).** Since we observed most of the differences between AdCa all (NCR + CR) versus NE, we proceeded with the rest of the analyses comparing NE vs AdCa all.

### The LuCaP PDX tumor proteome is consistent with established NE and AdCa gene signatures

To evaluate differences in the overall proteome landscape between NE (AR-) and AdCa (AR+) PDX samples, we performed an unsupervised clustering of all the proteins measured. We observed and confirmed the previous remarks that there is a distinct intra-and inter-tumor variability across all PDX samples **(Fig 2A).** More specifically, there is more variability across intact mice (NCR) and castrated (CR) AdCa PDX pairs than all of the NE samples, which indicates that the relative protein abundance in NE PDX samples are more similar than we might have expected. We then evaluated if this variability is consistent with the top 50 most highly upregulated proteins across all samples and at this level, the data showed that AdCa PDX tumors clustered uniquely and distinctly from the NE tumors **(Fig 2B).**

**Figure 2.**
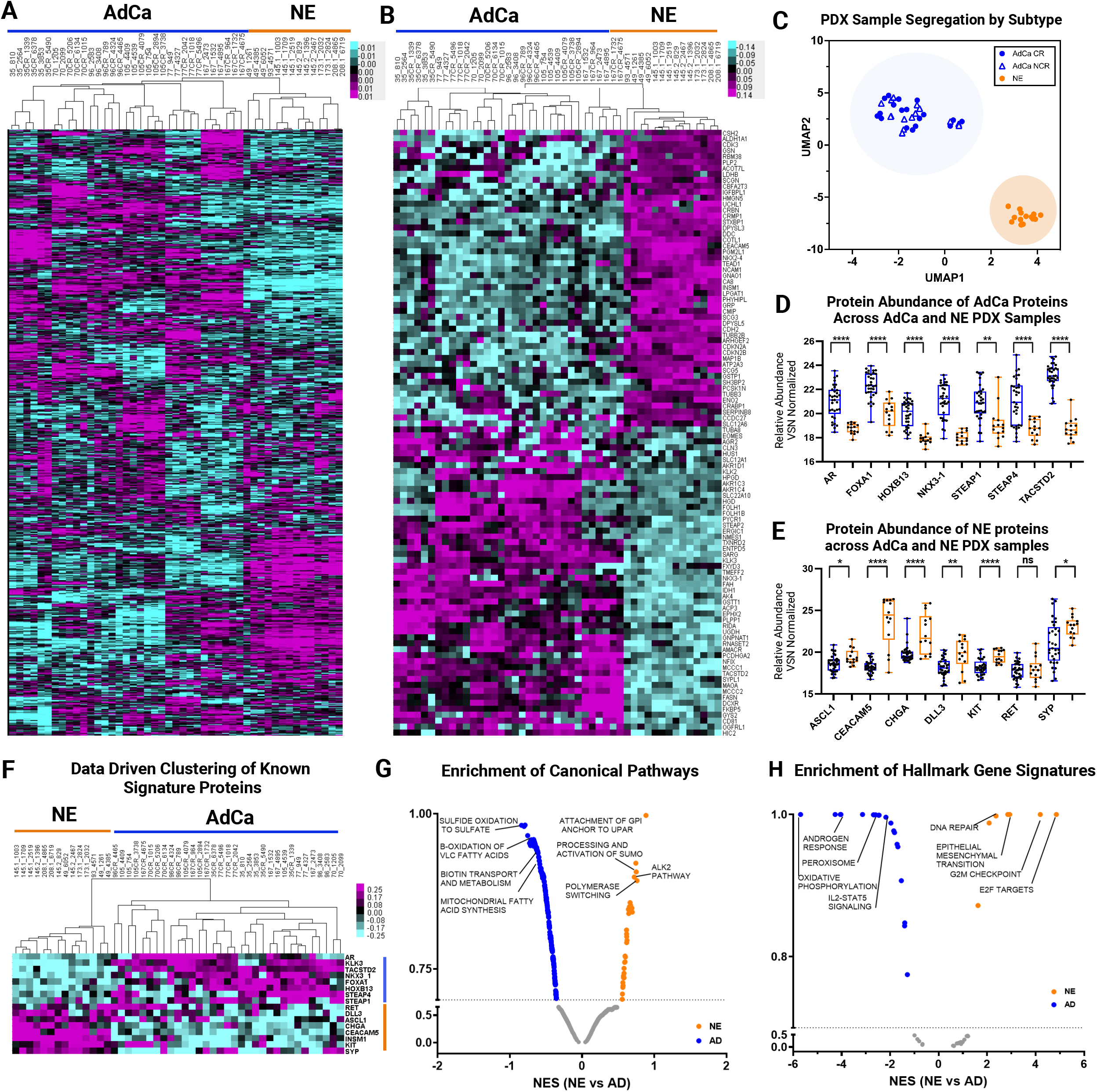
Proteomics landscape of PDXs in Prostate Cancer. **A.** Unsupervised clustering data drive 7738 master proteins 1% FDR. **B.** Unsupervised clustering of TOP 50 NE and 50 AdCA proteins. **C.** UMAP analysis of all PDXs. **D.** Data driven supervised hierarchical clustering of NE and AdCa signatures proteins. **E.** NE signature proteins**. F.** AdCa signature protein. **G.** Pathway Analysis of NE and AdCa highlighting the top 3 pathways on each group. **H.** Hallmarks in cancer analysis of NE and AdCa highlighting the top 4 pathways on each group (FDR 0.25).

Unsupervised data driven clustering and UMAP analysis showed that the PDX tumors clustered within their sub-groups despite the intra-and inter-PDX tumor variability (**Fig 2C**). Most importantly, individually analyzed proteins that have been shown to differentiate NEPC (ASCL1, RET, CEACAM5, CHGA, DLL3, SYP, KIT) vs CRPC AdCa (AR, FOXA1, HOXB13, NKX3-1, STEAP1, STEAP4, TACSTD2) showed that the proteins measured that are different in expression between NE vs AdCa are respectively hyper-abundant in their corresponding group comparisons (Fig D-F). Pathway analysis showed that there are 34 pathways enriched in NE and 267 in AdCa. Among others, pathways enriched in the AdCa PDX tumors consisted of sulfide oxidation to sulfate, β-oxidation of very long chain fatty acids (VLC-FA), (**Fig 2G**). In the NE PDX tumor samples, we identified an enrichment of proteins involved in the processing and activation of SUMO, ALK2 pathway, polymerase switching, and attachment of GPI anchor to u-PAR (urokinase-type plasminogen activator) (**Fig 2G**). Between both groups, these pathways are involved in commonly known hallmark gene signatures regulating proliferation, E2F targets, metastasis, adhesion, oxidative phosphorylation and angiogenesis through cell signaling (**Fig 2H**)and plays an important role in the tumor microenvironment ^36, 37^.These results provide evidence that the PDX tumor protein profiles maintain the proteome architecture and footprint similar to the clinical phenotypes previously observed.

### The LuCaP PDX tumor phosphoproteome reveals increased inter-patient homogeneity with established NE and AdCa gene signatures

To evaluate the overall phosphoproteome in the LuCaP PDX samples, we performed a sequential metal oxide phospho-enrichment targeting Serine (S), Threonine (T), and Tyrosine (Y) residues (**Fig 1A. II).** Unsupervised hierarchical clustering showed that the AdCa PDX samples LuCaP 96CR (replicate 10C), LuCaP 105, LuCaP 105CR, LuCaP 167 and LuCaP 167CR clustered more closely with 49 (replicate 3A), 145.2, 173.1 and 208.1 of NE origin and NE PDXs 49 and 145.1 clustered more closely to AdCa LuCaP 35, 35CR, 70, 70CR, 96, 96CR, 77, and 77CR **(Fig 3A).** This indicates that the phosphoproteome has more cross variability between NE and AdCa PDX tumors than the proteome, although the clustering patterns seems to reflect less interpatient heterogeneity when compared to the proteome. These data provide a new insight into the canonical understanding of these two mCRPC subgroups where the phosphorylation signatures might have more signaling overlap, allowing for the testing of novel drug targets that may treat both AdCa and NE tumors. When clustering the top 50 proteins that mapped to hyper-phosphorylated peptides from each of the AdCa and NE PDX tumors there was distinct segregation between those sub-groups **(Fig 3B).** Similar to the proteome, we assessed the known NE and AdCa proteins and mapped them to their corresponding phosphorylated peptides. Data driven analysis, strikingly showed that these phosphopeptides clustered similarly to the NE and AdCa proteins (**Fig 2D**) even though most of these phosphoresidues have never been analyzed in this context **(Fig 3C).** Since the unsupervised clustering showed that the NE and AdCa PDX tumor samples were not grouped based on pathological phenotype (**Fig 3A**), we evaluated their overall dimensionality using UMAPs that showed that these samples segregated within their corresponding sub-groups **(Fig 3D).** Using the proteins from the volcano plot (**Fig 1F)**, we mapped several functional phosphoproteins that have drugable phosphosites (i.e. phosphopeptides that map to phosphoresidues on proteins with known functional activity) that were unique in AdCa (AR_pS651, pS310, pS120; MYC_pT58; RB1_pT373 and SIRT1_pS47) and NE (STMN1_pS38; ADD2_pS697 and pS701; RET_pS696; CDK1_pT161; ARHGEF2_pS956, pS960; MCM2_pS108; EZH2_pT345; USP16_pS552; and E2A_pS379) PDX tumors **(Fig 3E).** Kinase Substrate Enrichment Analysis (KSEA) was performed to evaluate a proxy of kinase activity based on the phosphorylated substrates of the phospho-peptides measured. The results show that there are distinct kinases expressed at differential levels between NE versus AdCa PDX samples (**Fig 3F**). Furthermore, the phosphoproteome analysis showed that the hyperphosphorylated peptides enriched for pathways involved in metabolism of RNA, RNA processing, and PID-HIF TF pathway in AdCa; and chromatin modifying enzymes, neurexins and neuroligins, and transcription regulation by RUNX1 in NE PDX tumors **(Fig 3G)**. This confirms that NE PDX tumors are more closely related to a neuronal phenotype, while AdCa is more metabolically defined. As expected, gene set enrichment analysis (GSEA) indicated that androgen response, hypoxia and MYC targets were enriched in AdCa, while G2M checkpoint and E2F targets were enriched in the NE PDX tumors **(Fig 3H)**. Therefore, despite the global phosphoproteome clustering differences, there is a fundamental fidelity between AdCa and NE including drugable targets.

**Figure 3.**
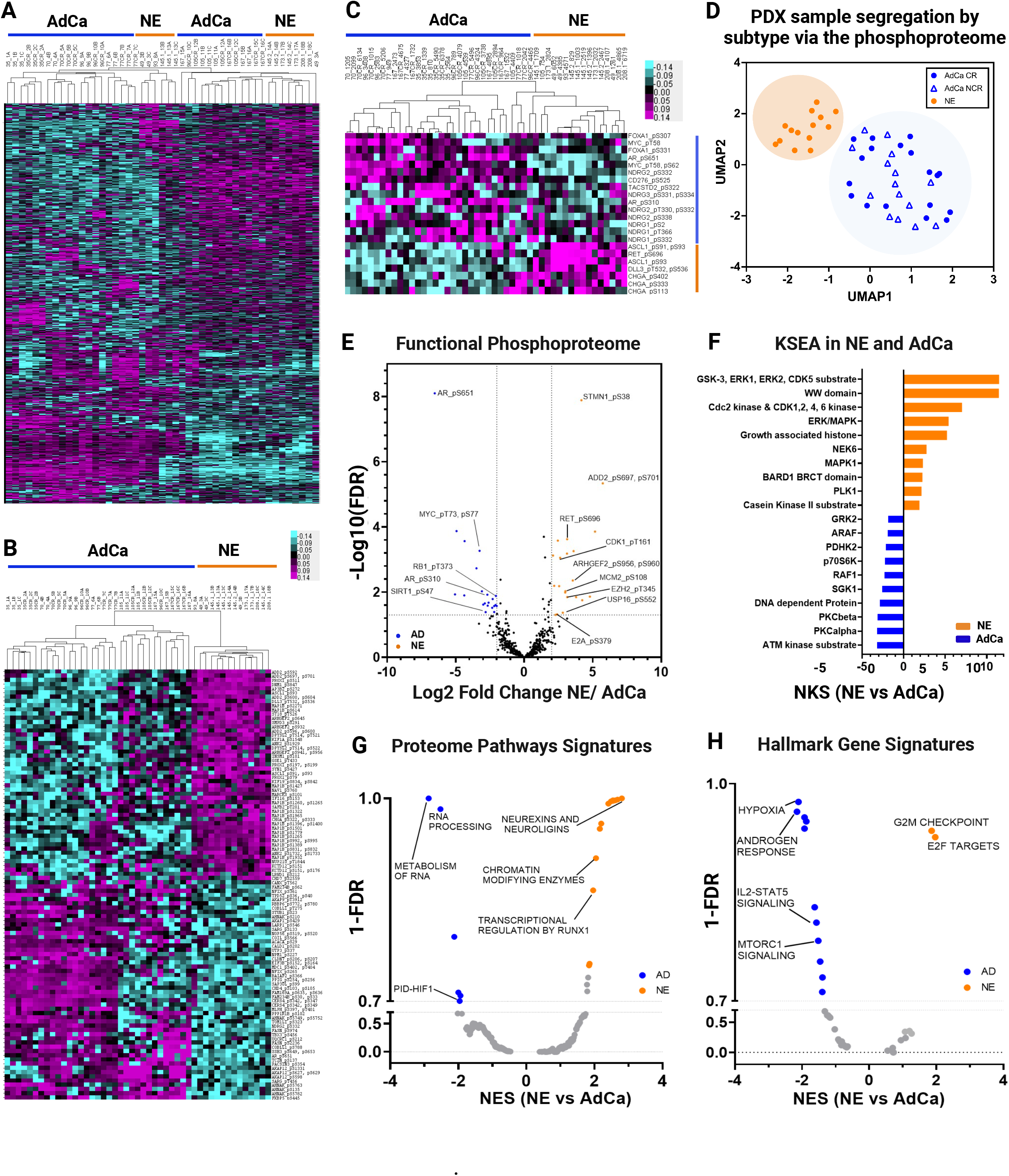
Phosphoproteomics landscape of PDXs in Prostate Cancer. **A**. Data driven unsupervised clustering of 9723 phosphopeptides with 1% FDR. **B.** Unsupervised clustering of TOP 50 NE and 50 AdCA hyper-phosphorylated peptides. **C.** Unsupervised hierarchical clustering signature protein-phospho-counterpart. **D.** UMAP analysis of all phosphopeptides. **E**. Functional phosphoproteome of NE and AD hyperphosphorylated peptides. **F.** Kinase/substrate enrichment (KSEA) analysis identified unique and known kinases that were predicted from the phosphoproteome (Top 10 hits are showed on each group). **G-H.** Gene set enrichment analysis (GSEA) was performed to identify canonical pathways (F) and hallmarks in cancer (G) enriched in NE (orange) and in AdCa (blue). NES, normalized enrichment score; orange, hyperphosphorylated in NE and blue hyperphosphorylated in AdCa.

### LuCaP PDX tumor proteomic and transcriptomic integration reveals dissonance between mRNA and protein targets

It has been established that mRNA expression has low to moderate correlation to protein expression with a 40-50% concordance ^38–40^, which might misguide potential nomination of novel targets if evaluated at the mRNA level only. Therefore, we conducted a concordance analysis between mRNA and protein abundance from the LuCaP PDX samples to evaluate potential discrepancy and nominate targets confidently for a clinical assay or biomarker development. The LuCaP PDX mRNA data ^41^ from a publicly available source was analyzed against the LuCaP PDX proteomic data collected in this manuscript. We analyzed and compared the proteins that were statistically significant and hyper-abundant (> 1.5-fold change) in the NE and AdCa PDX tumors, 336 and 360 proteins, respectively, with the corresponding matching gene transcripts that were statistically significant (matching proteins and gene transcripts). When plotted each proteins’ relative intensity based average quantification (iBAQ) log2 fold change against the correspondent transcript FPKM log2 fold change, the overall linear regression correlation was low with a statistical significance r^2^=0.2359 as expected **(Supplemental Fig 2A).** Next, we performed concordance analysis where we focused only on the hyper-abundant proteins and their matching mRNA expression level counterparts. While all the proteins analyzed here were statistically significant and hyper-abundant, we observed that only 54% (NE) and 59% (AdCa) of the matching proteins/gene transcripts were concordant (**C**; mRNA and protein are upregulated and hyper-abundant, respectively) while more than 40% in NE and 35% in AdCa proteins were non-concordant either discordant level I (**DC.I**; mRNA is not altered/changed significantly and protein is hyper-abundant) or discordant level II (**DC.II**; mRNA is significantly downregulated while the protein is hyper-abundant) (**Fig 4A**). These data strongly indicate that we are missing important drug targets and tumor biology within these two PDX sub-groups if we were to focus only on the mRNA. We then analyzed the directionality of the proteins versus the RNA counterpart within the subgroups and the overall dynamic range of mRNA expression was greater in the concordant group than the relative abundance in protein expression (**Fig 4B**) compared to the non-concordant groups (DC.I and DC.II). We then performed two sub analyses focusing on the AdCa and NE PDX tumors alone to show the overall distribution of the mRNA FPKM vs protein iBAQ fold changes **(Fig 4C and 4D)**. After evaluation of the protein class analysis **(Fig 4E),** we observed that NE hyper-abundant proteins which were classified as discordant Level I (DC.I) and discordant level II (DC.II) were mainly categorized as transcriptional regulators (such as NKX2-4, SMARCD1, ATF2, ZBTB21, MYEF2, and more), chromatin binding proteins (CENPH), and DNA metabolic proteins (TIPIN). This indicates that the mRNA transcripts of these proteins were not changed or the expression was downregulated while the protein was hyper-abundant **(Fig 4D).** These targets, have relevance in the biology of NEPC and would have been missed if only the mRNA transcripts were analyzed.

**Figure 4.**
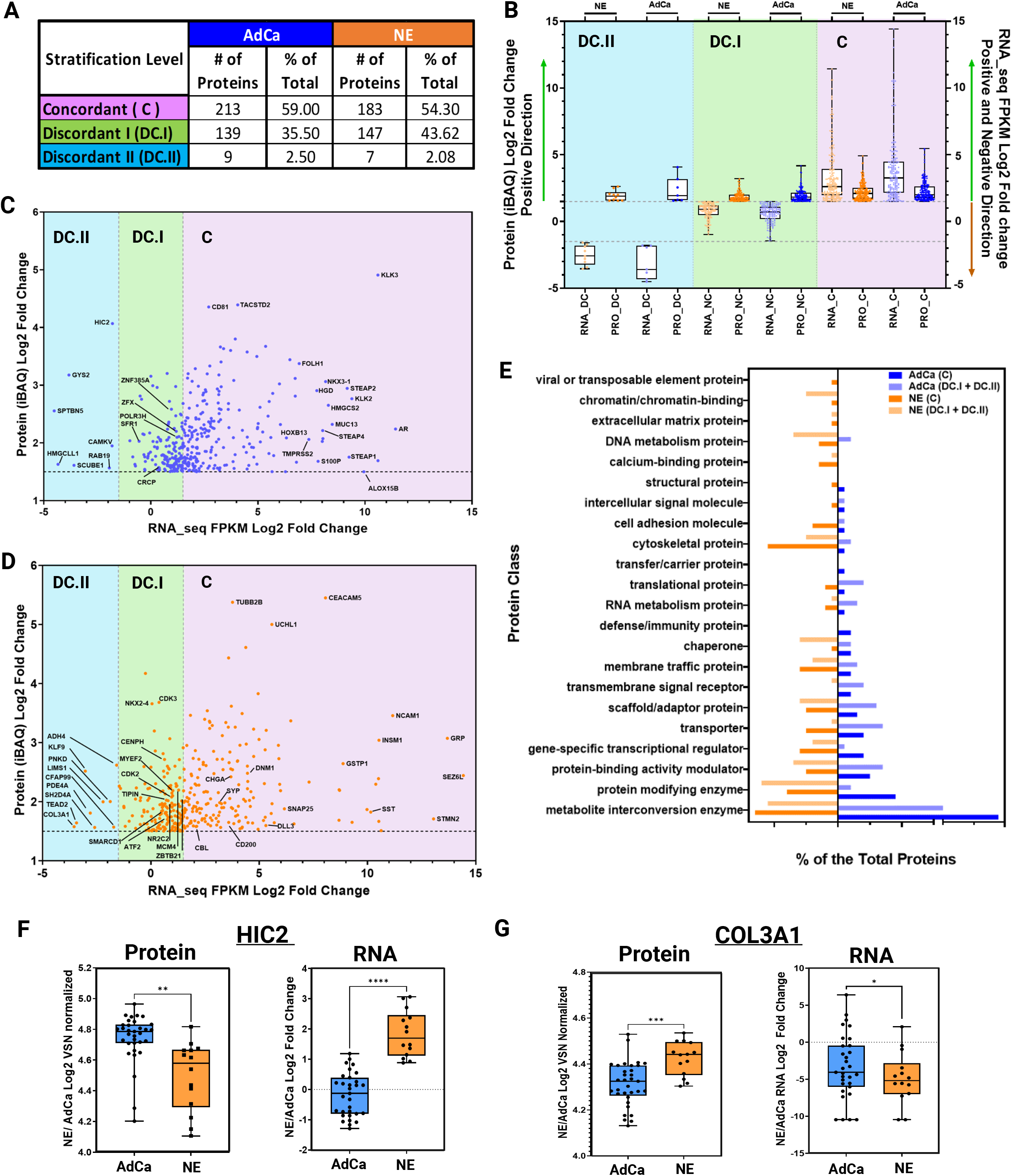
Proteomics and Transcriptomics data integration reveals functional overall dissonance and targetable proteins. **A.** Table shows the three main stratification levels of protein and RNA expression agreements, concordant (**C**); discordant I (**DC.I**); discordant II (**DC.II**) and the total number of hyper-abundant proteins in AdCa (n = 361) and in NE (n= 337) including percent distribution of total, respectively. **B.** Protein and RNA Log2 fold change evaluating only the hyper-abundant protein in NE (337 proteins) and AdCa (361 proteins) and simultaneously evaluating the direction of the mRNA expression of those proteins that are stratified as concordant (**C**; RNA and protein are upregulated and hyper-abundant), discordant I (**DC.I;** mRNA is not altered significantly and protein is hyper-abundant) and discordant II (**DC.II**; mRNA is significantly downregulated while the protein is hyper-abundant). **C-D.** AdCa and NE hyper-abundant proteins iBAQ (VSN normalized and ROTS p-value adjusted < 0.05 significances) RNA FPKM (ROTS normalized and p-value adjusted < 0.05) Log2 fold change highlighting proteins of interest. **E.** GO protein class analysis of the NE and AdCa concordant and non-concordant plus discordant proteins. Box plots of protein log 2-fold change VSN normalized and RNA log 2-fold change of n = 33 AdCa and n=14 NE evaluating the overall expression in **F.** HIC-2 and **G.** COL3A1. Data are represented as mean ± SEM and. ∗∗p < 0.01, ∗∗∗p < 0.001, two-tailed Welch-corrected.

To identify if the proteins that are discordant level I (DC.I) and discordant level II (DC.II) share any common characteristics, we performed a gene ontology protein class analysis **(Fig 4E).** From this, we identified that the proteins involved in chromatin binding, DNA metabolism, chaperon, and protein modifying enzymes were over-represented in the NE discordant level I and discordant level II groups (DC.I + DC.II) compared to AdCa; while translational proteins, transporter, scaffold proteins, and metabolite interconversion enzymes were greater in the AdCa DC.I and DC.II groups. This data also shows that there was greater dissonance in the NE DC.I and DC.II than in AdCa PDX tumors indicating that these targets would have been disregarded if only mRNA would have been analyzed.

To illustrate an example of extreme discordance, such as in DC.II, patterns between protein and RNA, we show two examples of proteins that are highly abundant in AdCa (HIC2, **Fig 4F**) and NE (COL3A1, **Fig 4G**) but the mRNA expression levels are very low. HIC ZBTB Transcriptional Repressor 2 (HIC2), a protein that enables protein C-terminus binding activity, predicted to be involved in regulation of transcription by RNA polymerase. In prostate cancer, HIC2 protein expression was shown to be increased in tumors compared to benign hyperplasia and normal tissue with a Gleason score greater than 7 and grade 3 ^42^. Collagen Type III Alpha 1 Chain (COL3A1) is a protein that is found in most soft connective tissues along with type I collagen ^43, 44^. It is involved in regulation of cortical development, and it is the major ligand of ADGRG1 in the developing brain. Moreover, COL3A1 binding to ADGRG1 inhibits neuronal migration and activates the RhoA pathway by coupling ADGRG1 to GNA13 and possibly GNA12 ^45^. In prostate cancer, COL3A1 is suppressed by miR-29b in DU145, and after treatment with the miR-29b inhibitor COL3A1 expression is increased as well as the cells invasiveness ^44^. Therefore, in the case of these two proteins, these would have been missed if only mRNA was used to analyze differential changes. However, HIC2 and COL3A1 do play a vital role in prostate cancer that could lead to potential development of therapeutic targets.

### Systematic analysis of the functional proteome and phosphoproteome for actionable targets

We further investigated if these hyper-abundant proteins and phosphoproteins from the NE and AdCa sub-groups have other functional characteristics that can provide avenues for biomarker and drug development. For these interrogations, we used the secretome (found in the extracellular matrix) ^46^ and blood proteins (found in blood plasma) from the Human Proteome Atlas, the surfaceome ^47^ and drug analysis from the Therapeutic Target Database ^48^, and Genomics of Drug Sensitivity in Cancer (GDSC) ^49^. We analyzed the proteins that were hyper-abundant (**Fig 5A, 5B**) and hyper-phosphorylated **(Fig 5C, 5D)** in AdCa and NE PDXs and searched them against the databases. For this, we included concordance stratification level on the proteome. For the phosphoproteome, concordance was only possible by mapping the master protein concordance level from the proteome to the phosphorylated peptide counterpart **(Fig 5C and 5D).** This analysis shows that of the 82 and 70 hyper-abundant proteins in AdCa and NE, respectively, 44% (36 proteins) and 36% (25 proteins) have drug targets in different stages of development, while the remaining proteins have not been investigated (or identified), which can be used as potential therapeutic targets. Furthermore, while we were able to identify proteins that are hyper-abundant with functional attributes (secreted, surface and blood protein) and that a therapeutic treatment is available, our results also identified new 36 in AdCa and 25 in NE potential targets for future therapeutic development. This clearly indicates that interrogating RNA expression alone, not only we are missing new insights in the biology of prostate cancer but that we are also missing proteins that can be useful for therapeutic treatment development.

**Figure 5.**
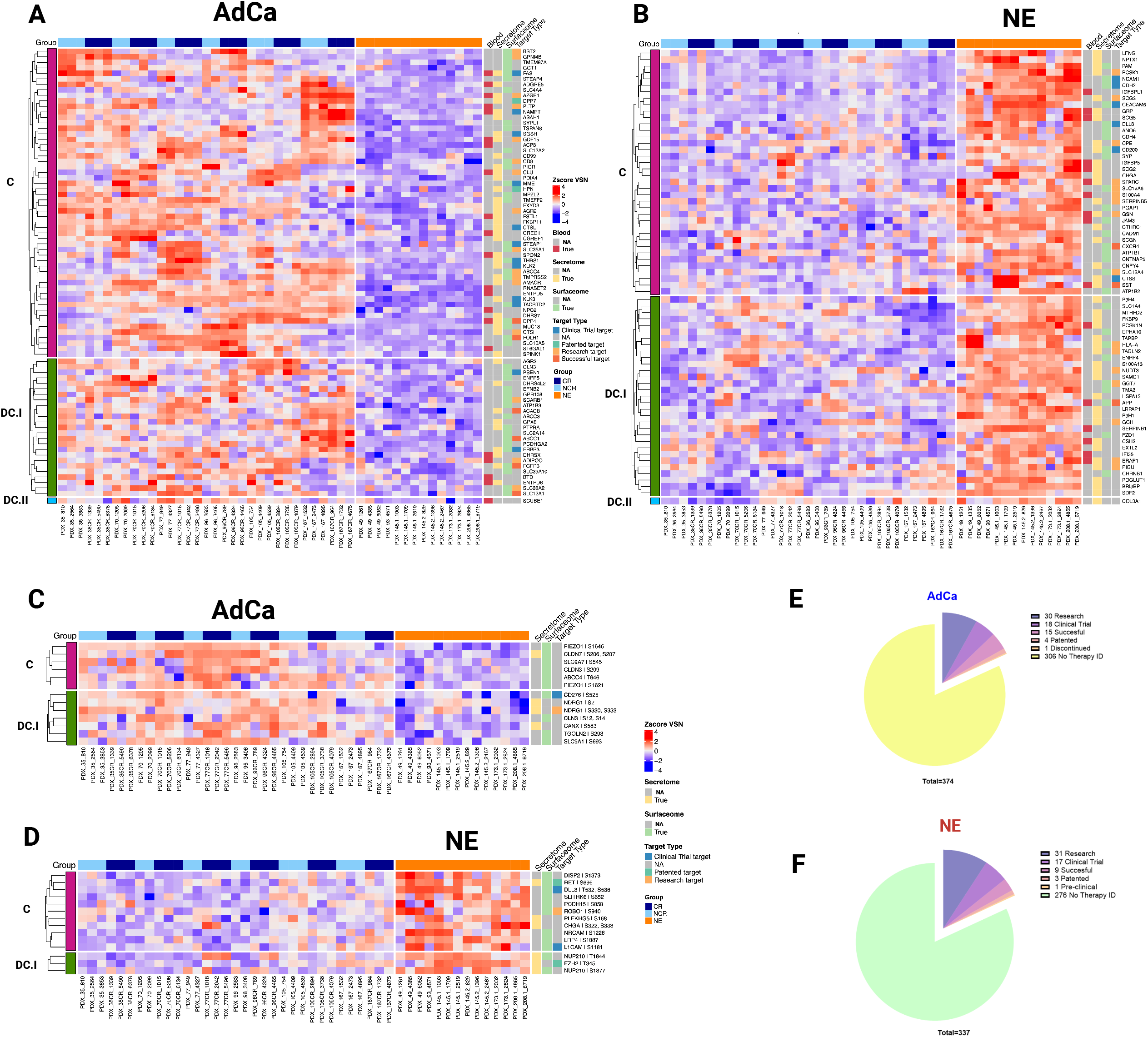
Functional proteome and phosphoproteome characterization. Heatmap data illustrates z-score VSN normalized protein hyper-abundance expression for AdCa (n=82) (**A.)** and NE (n=70) (**B.)** Heatmap data illustrates z-score VSN normalized proteins hyper-phosphorylated expression for AdCa (n=13) **(C.)** and NE (n= 14) **(D.)** Concordance level was defined by using the master protein counterpart and clustered based on these concordance from the proteome. Pie charts illustrate the therapeutic target distribution identified across the hyper-abundant proteins in AdCa **(E.)** with a total number 337 and NE **(F.)** with a total of 374 analyzed. Functional proteins color coding that are identified as Blood (red), Secreted (yellow) Surface (light green) and therapy target type such as clinical trial in blue, patented in green, research target in light orange, successful in terracotta and no available data identified as NA in gray.

At the phosphoproteome level, 16% (2 proteins) and 36% (5 proteins) hyperphosphorylated proteins in AdCa and NE, respectively, have drug therapies developed in different stages, while the remaining have not been investigated as potential therapeutic targets. Our data identified several proteins and phosphoproteins that have a localization characteristic viable for biomarker and potentially radio-ligand targeted therapy development while increasing our understanding of the prostate cancer biology. Overall, our data demonstrate that for certain targets there is a strong mRNA/protein concordance, but not for all, providing new insights in prostate cancer biology, identification and development of biomarkers, and drug targets.

## Discussion

The constant emergence of resistance to current treatment modalities in mCRPC requires identifying novel prognostic and predictive biomarkers and therapeutic targets for personalized patient stratification and development of new effective therapies.

We performed the first unbiased global proteomic and phosphoproteomics enrichment analysis of the LuCaP PDX models using the latest state of the art mass spectrometer, FAIMS as mentioned above ^31^. With this approach, we were able to measure over 8,600 master proteins, and demonstrated intra and inter PDX tumor variability. Furthermore, comparison of individual AdCa pair of intact mice (NCR) versus castrated (CR) tumors showed that their phenotypic proteome and phosphoproteome architecture maintains its fidelity with the known protein signatures and disease category; however, the differences between the pairs were not significant **(Fig 2A-D).** Interestingly, the unsupervised clustering analysis of the overall phosphoproteome between NE and AdCa revealed that neuroendocrine PDXs 49, 93, 145.1) clustered with AdCa PDXs, and AdCa PDXs (105, 105CR, 167, 167 CR and 96 CR-(replicate 10.C) clustered more closely with NE PDXs. However, when selecting the top phosphoproteins from NE and AdCa, the dichotomy of these two groups was more apparent indicating that most of the differences are present at the regulatory level of the protein such as phosphorylation. In addition, when we evaluated the phosphorylated AR, MYC, NDRG1, NDRG2 FOXA1, TACSTD2 in AdCa PDXs, and phosphorylated ASCL1, RET, DLL3, CHGA in NE PDXs, such as, these phosphopeptides clustered with their known protein/gene signature counterpart **(Fig 3D)**. The unsupervised clustering of these phosphorylated peptides was not previously characterized since phosphoprotein signatures have not been established or defined to date. However, we can clearly observe that these phosphoresidues do maintain their clustering profile with their respective disease phenotype. This is the first detailed analysis of phosphoproteome in different stages of prostate cancer and an outstanding resource for new therapeutic targets identification and increased insight of prostate cancer biology.

While genomes are typically constant, proteomes are quite different from cell to cell, both spatially and temporally. Proteins have over 400-post translational modifications, so the diversity generated from a single protein is larger than that of the corresponding gene ^50, 51^. Although all genes are present, not all genes are expressed in all cells, and moreover, proteins are expressed differently in different cell types depending on the tissue microenvironment. Many attempts have been made to correlate mRNA with proteins ^33, 39, 40^. This has uncovered that for a given amount of protein to be translated, it will depend on the gene classification that is transcribed such as a metabolic gene (required for survival) versus a transcription factor (will be turn on/off or degraded as needed for different biological responses), the current cell state, and the post-translational modification that is driving the signal. Therefore, prediction of protein abundance or activity based on mRNA transcript levels leads to poor mRNA/protein correlation and potential misleading biomarker and drug target discovery. There are some current algorithms that implement different variables (time and space), including protein isoforms, that could potentially increase the probability of mRNA/protein predictability, but these are still a way off from true implementation ^52^.

The global unbiased analysis of the proteome and phosphoproteome indicates that there is more inter variability across the PDX samples at the phosphoprotein level than at the proteome, clearly demonstrating that phosphorylated events are more significant to the biology than predicting protein functionality by genomics and transcriptomics alone. Unfortunately, use of phosphoprotein analyses is currently hindered by the ability to generate robust phosphoprotein data from small tissue amounts.

In addition to evaluating the overall proteome landscape of the LuCaP PDX tumors, we took a step further by integrating transcriptomic ^41^ data to the proteome. There are two main goals of this work; 1) to investigate concordance/discordance of RNA and protein expression, and 2) to identify protein-based biomarkers that can lead to development of therapeutic targets. Our focus was on identifying the functionality of these proteins as defined as secreted, found in blood, and expressed on the cell surface as these would be attractive and feasible targets. We initially performed a traditional spearman’s correlation that did not identify any significant targets due to small sample size (**Supplemental Figure 2A**). However, we were able to evaluate the overall dissonance between mRNA and proteins by comparing the directionality of the differential protein (focusing on the hyper-abundant) and mRNA (focusing on any directionality: upregulated, downregulated, or not changed) expression of NE vs AdCa PDXs. Interestingly, several of the proteins known to be involved in prostate cancer biology showed discordance between protein and mRNA levels, a finding that may have future clearly points out to shortcoming of using mRNA expression in clinical disease management implications.

NEPC (AR-) has been defined as a disease that is highly transcriptionally active regulating proteins involved in DNA replication (for example DNA polymerase, thymidine kinase, dihydrofolate reductase and cdc6), and chromosomal regulation ^53^ while AdCa (AR+) prostate cancer is highly metabolically ^54^ regulated. Furthermore, it has been shown that mRNA from transcription factors have increased average decay rates compared with other mRNA transcripts and are enriched for “fast-decaying” mRNA with a half-lives <2 hours ^38^. On the other hand, mRNAs related to biosynthetic proteins have decreased decay rates and are deficient in fast-decaying mRNAs ^38, 39^.

Therefore, the identification of discordance observed between RNA and protein expression levels, on transcription factors and transcriptionally regulated proteins in these two CRPC disease states (AR+ and AR-), might be explained mainly by the nature of the protein function on the cell such that transcriptionally regulated RNA is more dominantly affected than metabolically involved RNA, leading to more or less discordance with the protein counterpart. In addition, this phenomenon can be explained further by comparing half-lives between proteins and mRNA as well the timing of data collection. Also, there are proteins involved in RNA metabolism that would have been missed equally since these kinds of RNA molecules would have degraded much faster (2-10 hours) than their respective proteins (10-46 hours)^55^. Therefore, nominating biomarkers and subsequently designing clinical assays based on the most stable molecule, such as proteins, and evaluating if these proteins are either metabolically or transcriptional involved will be highly recommended for assay development decision making.

In conclusion, we generated the largest proteome and phosphoproteome resource database on clinically relevant and widely used CRPC LuCaP PDX models. Our analysis showed that the overall proteome maintained its fidelity with known CRPC AdCa (AR+) and NE (AR-) markers. We found proteins that are known to be overexpressed and hyper-phosphorylated such AR, RET, ASCL1, DLL3, KIT, CECAM, PSMA/FOHL1 and novel proteins specifically with important functional characteristics for biomarker or drug development, such as surface localization, secreted to the extracellular matrix and/or found in blood plasma. Furthermore, our analyses showed that there is discordance between multiple proteins and their RNA counterpart that is more dominantly found in transcriptional regulated proteins compared to metabolic proteins. Future follow up studies where both RNA and proteins are collected at the same time and measured in parallel, will be highly recommended to rule in/out any temporal changes that might affect the RNA levels to the protein expression. We encourage multi-omic level analysis and incorporating the proteome to be considered as a vital element for biomarker and drug development and for effective personalized medicine.

## Acknowledgements

**ZES** is supported by the Department of Defense Prostate Cancer Research Program E01 W81XWH-20-1-0070-P00002 PC190210JMD and Molecular, Genetic, and Cellular Targets of Cancer Training Program. The establishment and characterization of the LuCaP PDX models were supported by the Pacific Northwest Prostate Cancer SPORE P50 CA97186, the P01 NIH grant CA163227, and the Institute for Prostate Cancer Research. We would like to thank the Comparative Medicine Animal Caregivers for assistance with the LuCaP PDX maintenance and the patients and their families who generously donated tissues that made this research possible. JMD is supported by the NIH R01CA269801 and the Department of Defense Prostate Cancer Research Program W81XWH-18-1-0542. The content is solely the responsibility of the authors and does not necessarily represent the official views of the National Institutes of Health.

## Author Contributions

**ZES** executed, coordinated, and wrote all parts of the manuscript draft with support from **JMD**. **GL** helped process the PDX tumours for protein extraction and clean up. **JHH, HB, AD, AA, EB** helped with the proteome and phosphoproteome data normalization and analysis**. JHH** and **JMD** helped with conceptualization. Resources were provided by **EC, SRP** which includes the PDX LuCaP tumours and **ML** with scripts for proteome analysis. **PSN** and **IC** generated the RNA data from the PDX tumours. All authors contributed to the article and approved the submitted version. All authors provided constructive critiques in editing the manuscript.

## Inclusion and Diversity Statement

## Materials and Method

### LuCap Patient Derived Xenografts Tumors

A total of 48 PDX tumors from the LuCap series ^29^ representing 18 LuCap PDX models (2-3 independent tumors samples per model) were used for this analysis. These PDXs were obtained in a frozen pellet that originated from subcutaneously implanted patient-derived advanced prostate cancer from primary tumors and metastatic sites as described previously ^29^.

### Cell Prep Protein Extraction from PDX Tumors for Phosphopeptides and Proteome Enrichment

Approximately 150 mg of each tumor was processed as previously described ^32^; The lysis buffer contained 7M urea, 2M thiourea, 0.4M Tris pH 8.0, 20% acetonitrile (ACN), 10 mM TCEP, 25 mM chloroacetamide, Thermo Scientific’s Halt protease inhibitor cocktail 1x concentration (originally 100x), and phosphatase inhibitors (HALT from Thermo). After adding 500 uL of the lysis buffer to the tumor pieces, samples were placed on ice, then vortexed and centrifuged at 12000 x g for 10 min. The samples were then sonicated for 5 seconds using a probe sonicator set at 30% amplitude and kept on ice during the entire sonication process. After sonication, the samples were incubated for 0.5 hrs at 37°C, then at room temperature for 15 min to reduce and alkylate cysteines and centrifuged at 12,000 x g for 10 min at 18°C. Protein concentration was measured using Bradford Assay (Bio-Rad).

We then used 2.5 mg of protein, added 10 μg of 20 μg/mL Lysyl Endopeptidase (WAKO, 125-05061) and incubated the samples @ room temperature for 5-6 hr at pH 7.4. Then adjusted pH to 7.5-8.0 using 1 M Tris, pH 11. Then we added Worthington TPCK-treated trypsin (1 mg/mL) dissolved in 1 mM HCl supplemented with 20 mM of CaCl2 per to prevent autolysis. The trypsin mixture incubated at 4°C for about 1 hour prior to adding to the protein lysate. Samples were diluted 5-fold by adding 10 mM tris, pH 8.0 to dissolve urea <2M followed by trypsin addition at 1:50 trypsin/protein ratio overnight at 37°C.

After incubation, samples were acidified with TFA to pH 3 or less. Two sequential reverse phase extraction methods were used first. HLB was used first and then the flow thru from HLB and wash fractions were vacuum dried, resuspended, and cleaned up again using a C18 solid phase extraction method. The peptides were combined from HLB and C18 cleanups and peptide yield was measured using the BCA peptide assay (Thermo Fisher scientific cat# 23275). Samples digestion efficiency prior to mass spectrometry analysis was inspected by evaluating samples before and after enzymatic digestion using SDS-page gels and again after mass spectrometry analysis.

### Mass Spectrometry

1 ug of purified peptides was submitted for mass spectrometry analysis (Proteome) and 2 mg of peptides were kept and used for enrichment of phosphopeptides using the Sequential Metal Oxide Affinity Chromatography (SMOAC) Kit (Thermo Fisher Cat# A32993, A32992). The quantitative analysis of phosphoserine, phosphotyrosine and phosphothreonine Peptides by Quantitative Mass Spectrometry was performed as previously described ^56, 57^ with minor modifications in-tandem using SMOAC assay. The desalted peptide mixture was fractionated online using EASY-spray columns (25 cm 3 75 mm ID, PepMap RSLC C18 2 mm). The gradient was delivered by an easy-nano Liquid Chromatography 1000 ultra-high-pressure liquid chromatography (UHPLC) system (Thermo Scientific). Tandem mass spectrometry (MS/MS) spectra were collected on FAIMS TRIBRID mass spectrometer (Thermo Scientific). Samples were run in biological replicates, and raw MS files were analyzed using MaxQuant version 1.4.1.2 ^33^. MS/MS fragmentation spectra were searched using ANDROMEDA against the Uniprot human reference proteome database with canonical and isoform sequences (downloaded August 1^st^ 2021 from http://uniprot.org). N-terminal acetylation, oxidized methionine, and phosphorylated serine, threonine, or tyrosine were set as variable modifications, and carbamidomethyl cysteine (*C) was set as a fixed modification. The false discovery rate was set to 1% using a composite target-reversed decoy database search strategy. Group-specific parameters included max missed cleavages of two and label-free quantitation (LFQ) with an LFQ minimum ratio count of one. Global parameters included match between runs with a match time and alignment time window of 5 and 20 min, respectively, and match unidentified features selected.

### Mass Spectrometry Pre-processing of Proteomics and Phosphoproteomics Data

Maxquant imputed peptide level raw intensity files were obtained for each sample after the mass spectrometry experiments. We first summed the intensity of duplicated peptides based on the peptide sequences. In considering peptides with missed cleavages, we identified and then summed groups of peptides of variable length but had identical base sequences. We then mapped the peptide sequences to the most likely gene candidate based on algorithms by MaxQuant ^33^. We then averaged the intensity of the peptides that belonged to the same gene. To note, this averaging was only performed for the proteomics and not the phosphoproteomics data. At this stage, we aggregated all samples and then conducted VSN normalization for each dataset ^35, 58^. We omitted one 208.1 NEPC sample due to the unexpected expression of AR, which is normally only detected in ADCA. ROTS nomination of statistically significant features. To nominate statistically significant protein or phospho-peptides, we utilized ROTS ^59^. FDR adjusted p-values were computed in which we deemed less than 0.05 as statistically significant. This was used as the threshold for differentially represented features in ADCA or NEPC. We performed this for both proteomics and phosphoproteomics outputs. Hierarchical clustering was performed using the Cluster 3.0 program with the Pearson correlation and pairwise complete linkage analysis (Eisen et al., 1998). Java TreeView was used to visualize clustering results (Saldanha, 2004). Quantitative data for each phosphopeptide can be found in supplemental Table 5.

### Proteomics and Transcriptomics Correlation

RNA data was obtained for the same PDX samples ^41^ in which we averaged, by sample ID, the log2 FPKM data for the RNA sequencing or the VSN normalized protein abundance data. We only included genes that were detected in both datasets and then conducted Spearman’s correlations between the RNA expression and protein abundance levels for each gene across all samples. Concordance analysis was done by comparing the hyperabundant proteins from the NE and AdCa groups with the RNA levels in the transcriptomics data. Concordance indicates that the protein and RNA fold change is greater than 1.5. Non-concordance was defined as proteins that are greater than 1.5-fold change, with RNA between -1.5 to 1.5 fold change. Discordance is defined as proteins that are 1.5-fold change hyper abundant, with RNA is lower than -1.5 fold change. All were statistically significant with an adjusted p-value <.05.

### Protein Annotations and Databases

We analyzed protein functional annotations using Human Proteome Atlas (HPA version 22.0 http://www.proteinatlas.org/) that has categorized blood proteins, secretome ^46^ and surfaceome ^47^. Ontologies were identified using Gene Set Enrichment Analysis (GSEA version 4.2.1). Potential drug targets were further mapped to gene symbol, to PhosphoSitePlus ^60^, Therapeutic Target Database (TTD) ^48^ Genomics of Drug Sensitivity in Cancer (GDSC) ^49^, and HPA to acquire additional information on whether the targets had drug response data, or they were receptors, kinases, or known cancer/FDA-approved/potential drug targets.

## Statistical Analysis

All statistical data were presented after either t-tests or ROTS as described in the figure legends.

## Accession Numbers

LuCaP PDX RNA sequencing data is available at the Gene Expression Omnibus (GEO) under accession GSE199596. LuCap PDX phosphoproteome data is available at ProteomeXchange PDX042859 LuCap PDX proteome data is available at ProteomeXchange PXD042867

