## Supplemental Figures for "Unraveling the Global Proteome and Phosphoproteome of Prostate Cancer Patient-Derived Xenografts"

**S1.A**

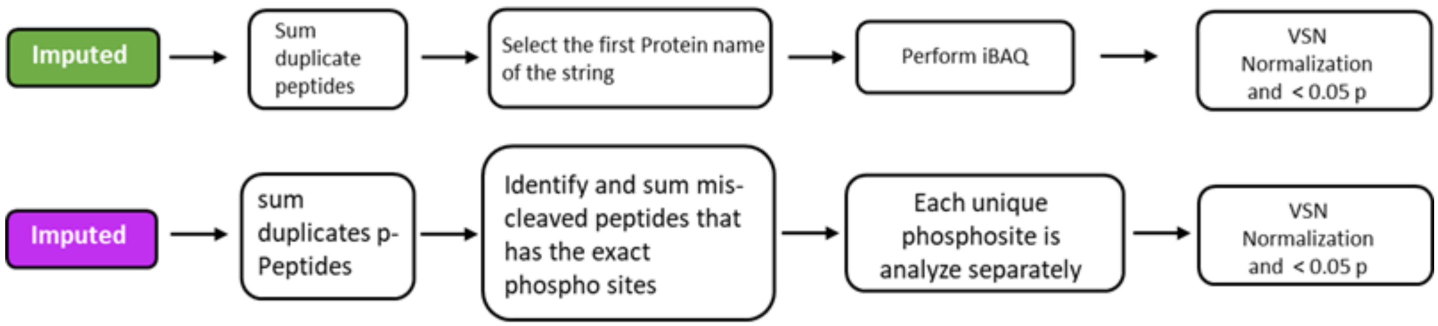


Sample processing workflow after raw files were searched using MaxQuant spectra matching to a peptide sequence, for the proteome (green) and for the phosphoproteome (magenta).

**S2.A**

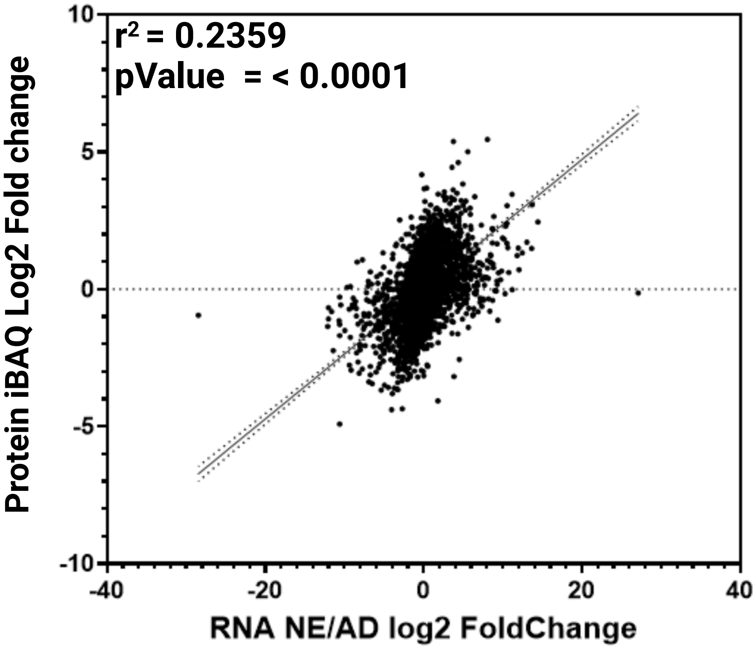


Overall linear regression plot between protein Log2 fold change relative intensity based average quantification (iBAQ) against the correspondent transcript FPKM Log2 fold change with a correlation of $r^2$=0.2359 and a p-value of < 0.0001
